# SLAMseq reveals transfer of RNA from liver to kidney in the mouse

**DOI:** 10.1101/2024.05.16.594511

**Authors:** Robert W Hunter, Jialin Sun, Trecia Palmer, Matthew A Bailey, Neeraj Dhaun, Amy Buck, James W Dear

**Affiliations:** Edinburgh Kidney Research Group, Centre for Cardiovascular Science, Queen’s Medical Research Institute, University of Edinburgh, Edinburgh Bioquarter, Edinburgh EH16 4TJ; Department of Renal Medicine, Royal Infirmary of Edinburgh, Edinburgh Bioquarter, Edinburgh EH16 4SA; Department of Pharmacy, the Affiliated Hospital of Qingdao University, Qingdao 266003, China; Centre for Immunity, Infection and Evolution, University of Edinburgh, Ashworth Laboratories, Charlotte Auerbach Road, Edinburgh EH9 3FL

**Keywords:** Extracellular RNA, exRNA, SLAMseq, liver injury

## Abstract

Extracellular RNA (exRNA) mediates intercellular communication in plants and lower animals; whether it serves a signalling function in mammals is controversial. Reductionist experiments, in which a single RNA is over-expressed or tagged, have shown RNA transfer between tissues but these may not be relevant to normal physiology. For example, the microRNA miR-122 is released from injured hepatocytes and is taken up by kidney cells. We sought to determine the scale of RNA transfer between liver and kidney through the metabolic labelling of RNA in mice. We used 4-thiouracil to specifically label RNA in hepatocytes then detected labelled (thiolated) RNA in the kidney using SLAMseq: SH-Linked Alkylation for the Metabolic sequencing of RNA. In the kidney, mRNA labelling was detected in 5% of all kidney transcripts under healthy conditions and was increased to 34% of kidney transcripts after acute hepatocellular injury. Labelling was evident in kidney transcripts mapping to known hepatocyte marker genes, to a greater extent than those mapping to markers of other cell types. Labelled small RNA was not detected in kidney tissue. Our results are consistent with the transfer of RNA from liver to kidney; this transfer is augmented in liver injury.

## Introduction

Extracellular RNA (exRNA) is an important mediator of intercellular communication in plants and lower animals. Small, non-coding RNAs travel between cells – even between organisms – to induce wide-ranging changes in gene expression *via* RNA interference. This phenomenon participates in anti-viral responses and regulates various facets of development and homeostasis.^1–5^

In mammals, although exRNA is found ubiquitously in biofluids, its role in intercellular signalling is controversial and lacks quantitative evidence.^6^ We do not know whether exRNA regulates physiologically important processes in higher organisms. Several features of mammalian exRNA biology are compatible with its serving an evolutionarily conserved signalling function. exRNA is protected from degradation by being packed into vesicular, lipoprotein and ribonucleoprotein carriers.^7,8^ exRNA transport bears many hallmarks of a physiological signalling system: cellular RNA export is often selective and uptake by recipient cells may be selective^9,10^ and regulated^11,12^.

In experimental rodent models, exRNAs can induce physiologically relevant changes in recipient cells when single microRNAs (miRNAs) are over-expressed, tagged or blocked. For example, adipose-derived miRNAs regulate Fgf21 expression in hepatocytes^13^ and bone-marrow-derived miR-155 regulates insulin metabolism in hepatocytes and adipocytes^14^. Similarly, there is evidence that miR-122 appears moves from the liver to the kidney in response to liver and systemic injury. In lipopolysaccharide-induced systemic inflammation, Rivkin *et al.* found elevated levels of mature miR-122 in the kidney, but not pre-miR-122, suggesting that it was not locally synthesised.^15^ We subsequently tested the hypothesis that liver-derived miR-122 was travelling to the kidney in the context of acute paracetamol overdose. After paracetamol-induced liver injury, miR-122 levels rose within kidney tissue; this effect was abolished after liver miRNA biogenesis was attenuated by hepatocyte-specific Dicer knockdown.^16^ It is important to seek a better understanding of how RNA signalling might mediate liver-to-kidney crosstalk because there is a poorly-understood association between liver and kidney disease.^17,18^

Such experiments perturbing single miRNAs provide compelling evidence that exRNA *can in principle* mediated intercellular signalling.^13,14,19–23^ Nevertheless, they are unable to resolve whether such signalling occurs – or is relevant – in a complex, physiological setting. It is likely that any intercellular signalling would involve multiple different RNAs, rather than a single dominant miRNA.

Therefore here, we sought to determine the global pool of RNA molecules moving between organs in the mouse using metabolic RNA labelling. We labelled hepatocellular RNA *in vivo* to test the hypothesis that RNA travels from the liver to the kidney. We determined the global pool of RNA molecules that follow this path in health and after acute hepatocellular injury.

## Results

### AAV8-TBG-Cre induces hepatocyte-specific expression of uracil phosphoribosyl transferase

We labelled RNA in hepatocytes using the modified nucleoside, 4-thiouridine. We sought to detect and sequence this labelled RNA within the kidney using SLAMseq: Thiol (<u>S</u>H)-<u>L</u>inked <u>A</u>lkylation for the <u>M</u>etabolic sequencing of RNA. To facilitate hepatocyte-specific RNA labelling, the adenoviral vector AAV8-TBG-Cre was used to induce hepatocyte-specific Cre expression in the floxed-stop-UPRT mouse. This is expected to remove a floxed stop cassette, turning on expression of recombinant HA-tagged uracil phosphoribosyl transferase (HA-UPRT).

We used four complementary methods to verify that this induced hepatocyte-specific expression of HA-UPRT, and in particular did not cause off-target UPRT expression in the kidney. In genomic DNA (gDNA), Cre-mediated recombination at the UPRT locus was detected within liver but not kidney, spleen or heart (Fig 1A; supplemental Fig S2A). Using podocin-UPRT mice as a positive control, we determined that this gDNA recombination assay was sensitive enough to detect recombination events occurring in as few as ∼1:500,000 kidney cells (Fig S2B). In RNAseq data, TBG promoter sequence was more abundant in liver after AAV8-TBG-Cre delivery but was no different in kidney (Fig 1B). When AAV8-TBG-Cre was injected into mTmG fluorescent reporter mice, green fluorescence was detected in hepatocytes but not in the kidney (Fig 1C). At the protein level, HA-UPRT expression was detected in liver after AAV8-TBG-Cre injection but not in kidney (Fig 1D).

**Fig 1.**
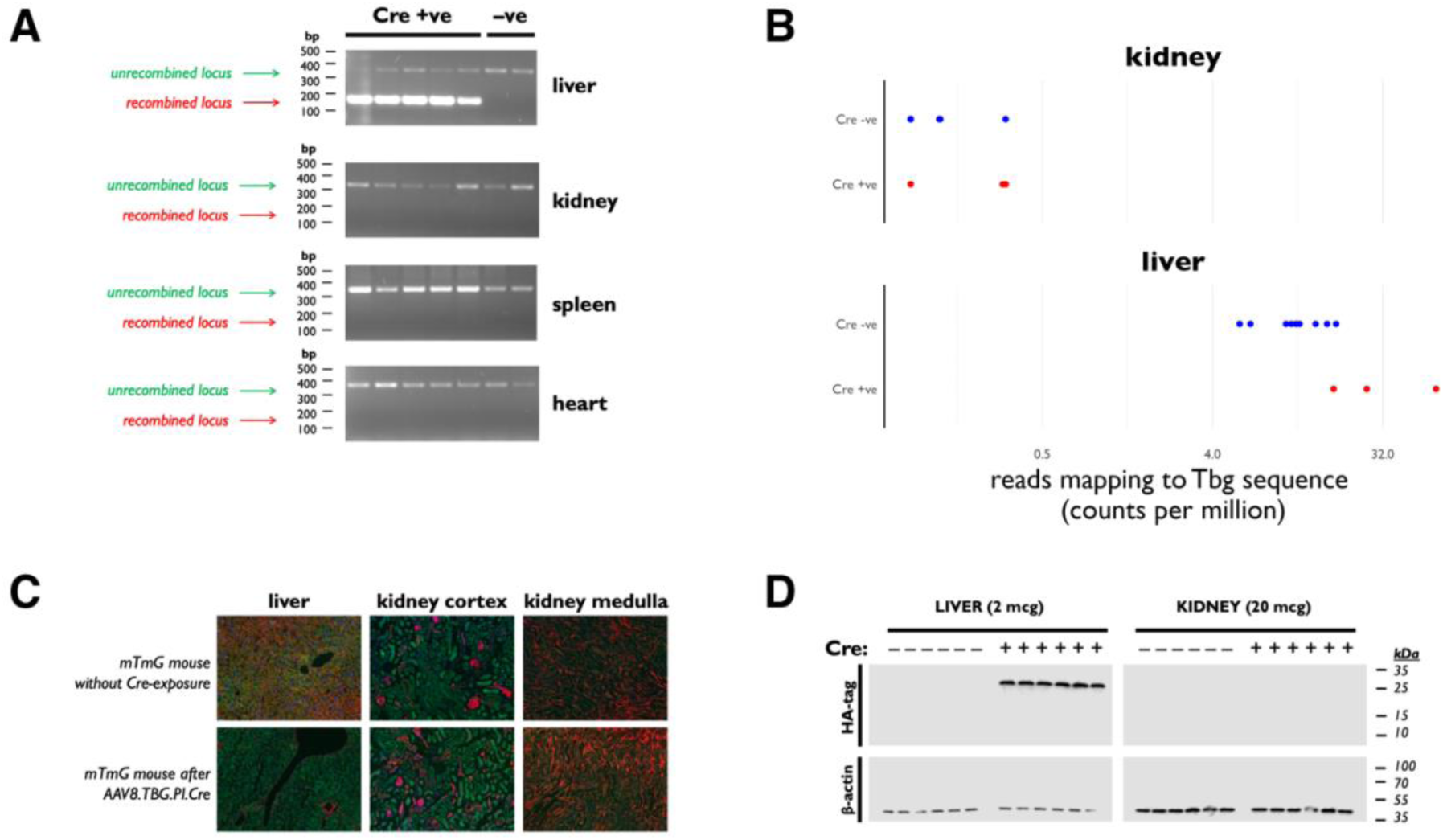
UPRT expression is induced specifically in hepatocytes. **A)** *Genomic DNA (gDNA) recombination assay.* gDNA was extracted from liver, kidney, heart and spleen and used as templates in a PCR assay designed to detect Cre-mediated recombination at the flox-stop-UPRT locus. The design of this assay is shown in supplemental FigS2A. Cre-mediated recombination was detected in the liver but not in kidney, spleen or heart. **B)***TBG promoter expression.* The AAV8-Cre vector contains the TBG promoter. In the RNAseq data obtained in the SLAMseq experiments, these sequences were over-represented only in liver samples, following AAV8-Cre injection. **C)** *Fluorescent reporting of AAV8-Cre activity.* The AAV8-Cre vector used to induce Cre recombination was injected into the mTmG fluorescent reporter mouse. Green fluorescence, indicative of Cre recombination, was detected only in hepatocytes and not in the kidney. **D)** *UPRT immunoblot.* Recombinant UPRT was detected by immunoblot (probing for the HA tag) in liver (2 micrograms of tissue homogenate per lane) and kidney (20 micrograms). β-actin was probed as a loading control. Cre-mediated UPRT expression was detected in liver but not kidney.

### 4TU labels hepatocyte RNA

We conducted an initial experiment in small groups of male and female mice (supplemental Fig S1A; Table S1). The purpose of this initial experiment was to verify that T>C conversions could be detected in liver RNA and to determine which control groups were necessary for the subsequent, larger, liver injury experiment. Four weeks after AAV8-TBG-Cre injection, mice were injected with 4TU. To allow sufficient time for 4TU to be incorporated into nascent liver RNA and then transferred to distant organs, we administered 4TU for 36 hours prior to organ harvesting. In a biotinylation assay, we determined that our protocol led to the successful incorporation of the thiol label into liver RNA (Fig 2A).

**Fig 2.**
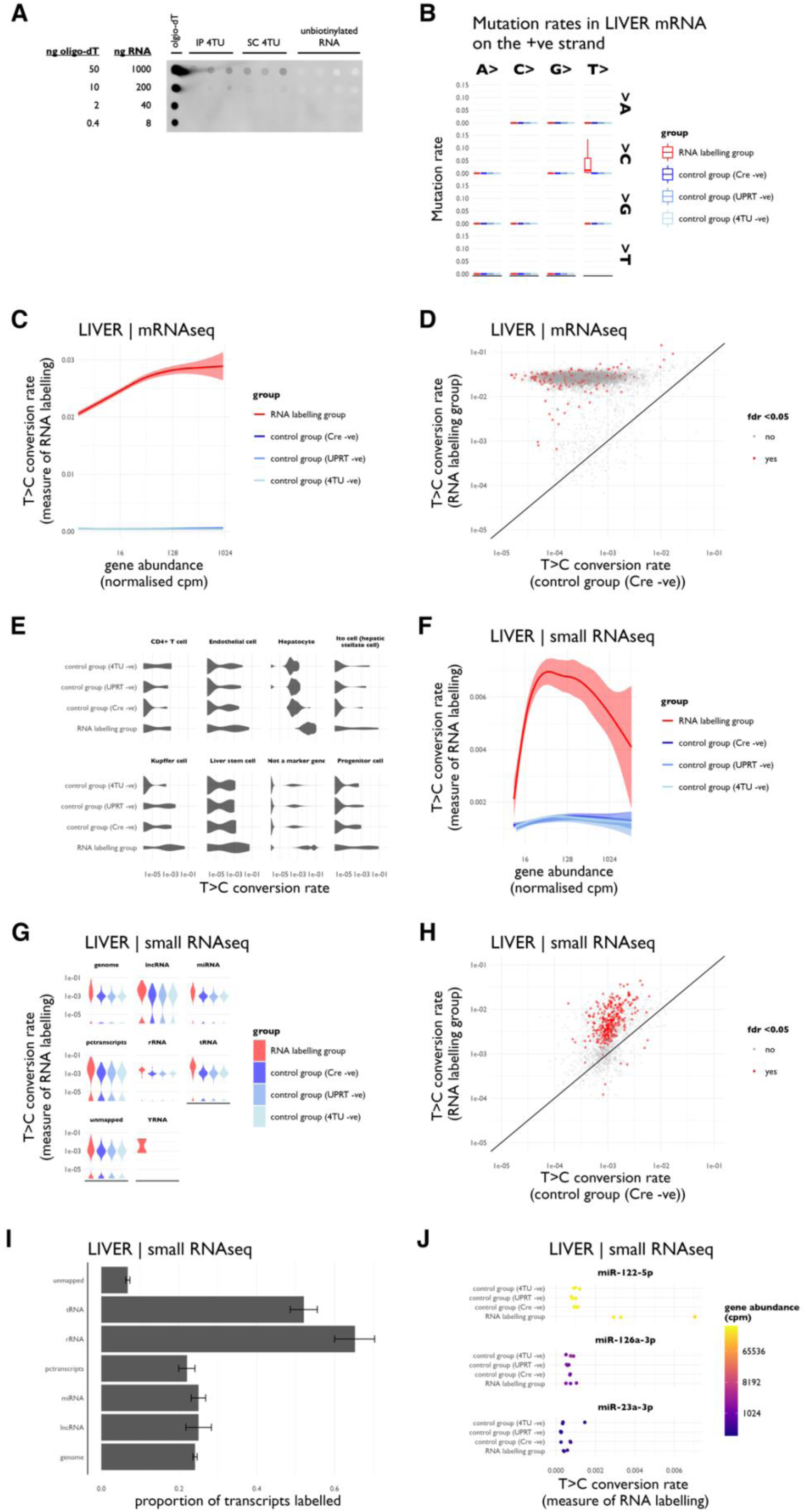
SLAMseq labels RNA in the liver in a small initial experiment. **A)** *Biotinylation dotblot.* Flox-stop-UPRT mice were treated with AAV8-Tbg-Cre and then injected with 4-thiouracil as either intraperitoneal or subcutaneous injections (n = 3 biological replicates in each group). Incorporation of the 4TU label into liver RNA was assessed using a biotinylation assay. RNA samples were biotinylated and then spotted in a dilution series next to a positive control (biotinylated oligo-dT) onto a nylon membrane. The membrane was probed with a Streptavidin-HRP conjugate. **B)** *Mutation rates in SLAMseq data.* Liver RNA was alkylated and sequenced in a SLAMseq protocol designed to analyse mRNA. Increased rates of T>C conversion were detected on the positive strand; increased rates of A>G conversion were detected on the negative strand (data not shown). **C)** *T>C conversion rates in mRNA SLAMseq data.* T>C conversion rate in liver mRNA. **D)** *Labelled RNAs in SLAMseq data.* Labelled RNA transcripts were detected by comparing gene-wise T>C conversion rates between Cre-positive (RNA labelling) and Cre-negative (control) groups using the beta-binomial method and setting a false discovery rate of 0.05. Each point represents a single gene mRNA; genes for which there was a significant between-group difference in T>C conversion rate are shown in red. **E)** *T>C conversion rates in known hepatocyte marker genes.* T>C conversion rates were determined in a pre-specified set of “marker genes”, known to be specifically expressed in defined cell types in other datasets. The marker genes are listed in supplemental Table S11. **F)** *T>C conversion rates in small RNA SLAMseq data.* Liver RNA was alkylated and sequenced in a SLAMseq protocol designed to sequence small RNA. Increased rates of T>C conversion were detected on the positive strand; increased rates of A>G conversion were detected on the negative strand (data not shown). The rates of T>C conversion are shown. **G)** *T>C conversion rates stratified by RNA biotype.* Increased rates of T>C conversion int the RNA labelling group were observed in reads mapping to all small RNA biotypes. **H)** *Labelled RNAs in SLAMseq data.* Labelled RNA transcripts were detected by comparing gene-wise T>C conversion rates between Cre-positive (RNA labelling) and Cre-negative (control) groups using the beta-binomial method and setting a false discovery rate of 0.05. **I)** *Labelling of small RNA in liver.* Stratified by RNA biotype. **J)** *miR-122, miR-126a and miR-23a labelling.* T>C conversion rates in the known hepatocyte-enriched miRNA, miR-122, and the known endothelial-enriched miRNAs, miR-126a and miR-23a. Data derived from male and female mice, n = 3 in each experimental group.

RNA was extracted from liver and kidney and prepared for SLAMseq. We first evaluated 4TU incorporation into liver mRNA. As expected, T>C conversions were observed in the SLAMseq dataset; rates of other single nucleotide conversion were unaffected (Fig 2B). T>C conversions were observed in the RNA labelling group at a rate that exceeded the baseline rate in Cre-negative, UPRT-negative and 4TU-negative control groups (Fig 2C; supplemental Table S3). The T>C conversion rate was higher in more abundant transcripts, consistent with more abundant RNA species being transcribed at a higher rate (Fig 2C). mRNA transcripts that exhibited significantly higher rates of T>C conversion in the RNA labelling group (compared to the Cre-negative control group) were identified using a beta-binomial test (Fig 2D; supplemental Tables S4 & S5). RNA labelling was apparent in transcripts known to be enriched in hepatocytes (Fig 2E).

We also used SLAMseq to assess labelling of small RNAs in the liver. An excess of T>C conversion was observed within the small RNA SLAMseq dataset (Fig 2F-H; supplemental Tables S4 & S6), although T>C conversion rates were markedly lower than in the mRNA dataset. RNA labelling was detected in reads mapping to all small RNA biotypes (Fig 2I). RNA labelling was evident in miRNAs known to be enriched in hepatocytes, including miR-122, whereas it was not evident in the known endothelial-enriched miRNA, miR-126a (Fig 2J).

### Labelled RNA is detected in the kidney

We next used SLAMseq to look for labelled RNA in the kidney after hepatocyte-specific RNA labelling. We detected a small excess of T>C conversions in kidney mRNA (p < 10^−9^; Fig 3A-C; supplemental Table S3). In a gene-wise analysis, 0.6% of all mRNAs in the kidney were identified as having been labelled (supplemental Table S4). In contrast, there was no increase in T>C conversion rate in kidney small RNA (Fig 3D-E).

**Fig 3.**
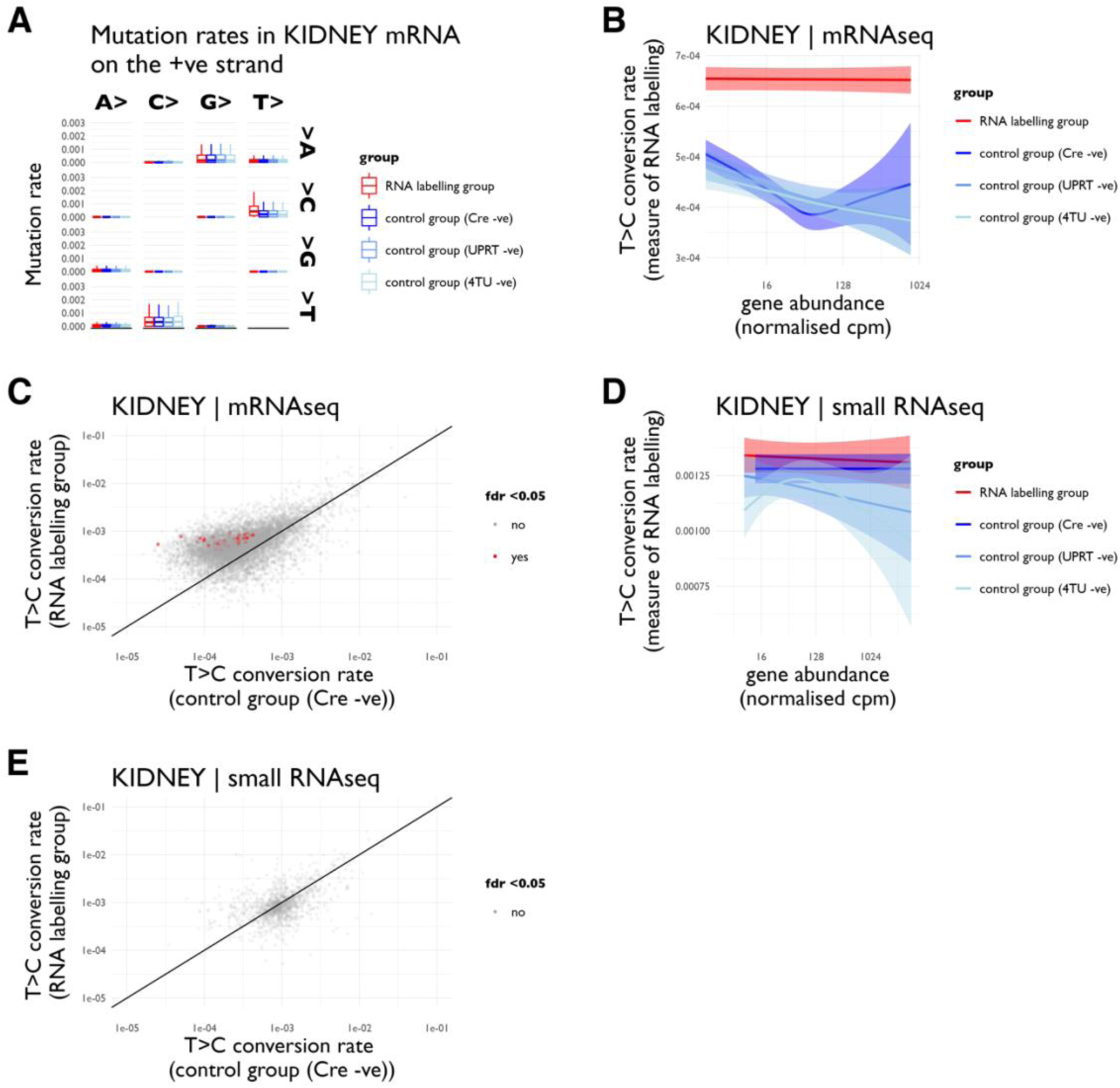
Labelled RNA is detected within kidney in a small initial experiment. **A)** *Mutation rates in SLAMseq mRNA data.* Kidney RNA was alkylated and sequenced in a SLAMseq protocol designed to sequence mRNA. Increased rates of T>C conversion were detected on the positive strand; increased rates of A>G conversion (not shown) were detected on the negative strand. **B)** *T>C conversion rates in SLAMseq mRNA data.* T>C conversion rate in kidney mRNA. **C)** *Labelled RNAs in mRNA SLAMseq data.* Labelled mRNA transcripts were detected by comparing gene-wise T>C conversion rates between Cre-positive (RNA labelling) and Cre-negative (control) groups using the beta-binomial method and setting a false discovery rate of 0.05. **D)** *T>C conversion rates in small RNA SLAMseq data.* **E)** *Labelled RNAs in small RNA data.* In contrast to the mRNA data, no small RNA labelling was detected within the kidney. Data derived from male and female mice, n = 3 in each experimental group.

### 4TU labels hepatocyte RNA after acute hepatocellular injury

To examine how liver-to-kidney RNA transfer might change in the context of liver injury, we induced an acute hepatocellular injury by paracetamol overdose (supplemental Fig S1B; Table S2). To reduce variation in the injury response, we performed these experiments only in male mice (n = 6), alongside a group of male mice subjected to RNA labelling under healthy conditions (4TU +ve, AAV8-TBG-Cre +ve, paracetamol −ve; n = 6) and male negative control groups (4TU +ve, AAV8-TBG-Cre −ve; n = 6 and 4TU −ve; n = 3). As the DMSO vehicle ameliorated paracetamol-induced liver injury when delivered intraperitoneally, we used a subcutaneous route for 4TU in these liver injury studies. We confirmed that the subcutaneous route induced RNA labelling to the same extent as intraperitoneal 4TU (Fig 2A).

A single dose of intraperitoneal paracetamol (300 mg/kg) induced a sublethal, acute hepatocellular injury with extensive necrosis (Fig 4A-D). mRNA from liver exhibited global transcriptional changes compared to control groups (Fig 4E&F); differentially regulated genes were enriched in GO and KEGG terms relating to inflammation and repair, as well as homeostatic hepatocyte functions such as lipid and xenobiotic metabolism (Fig 4G&H).

**Fig 4.**
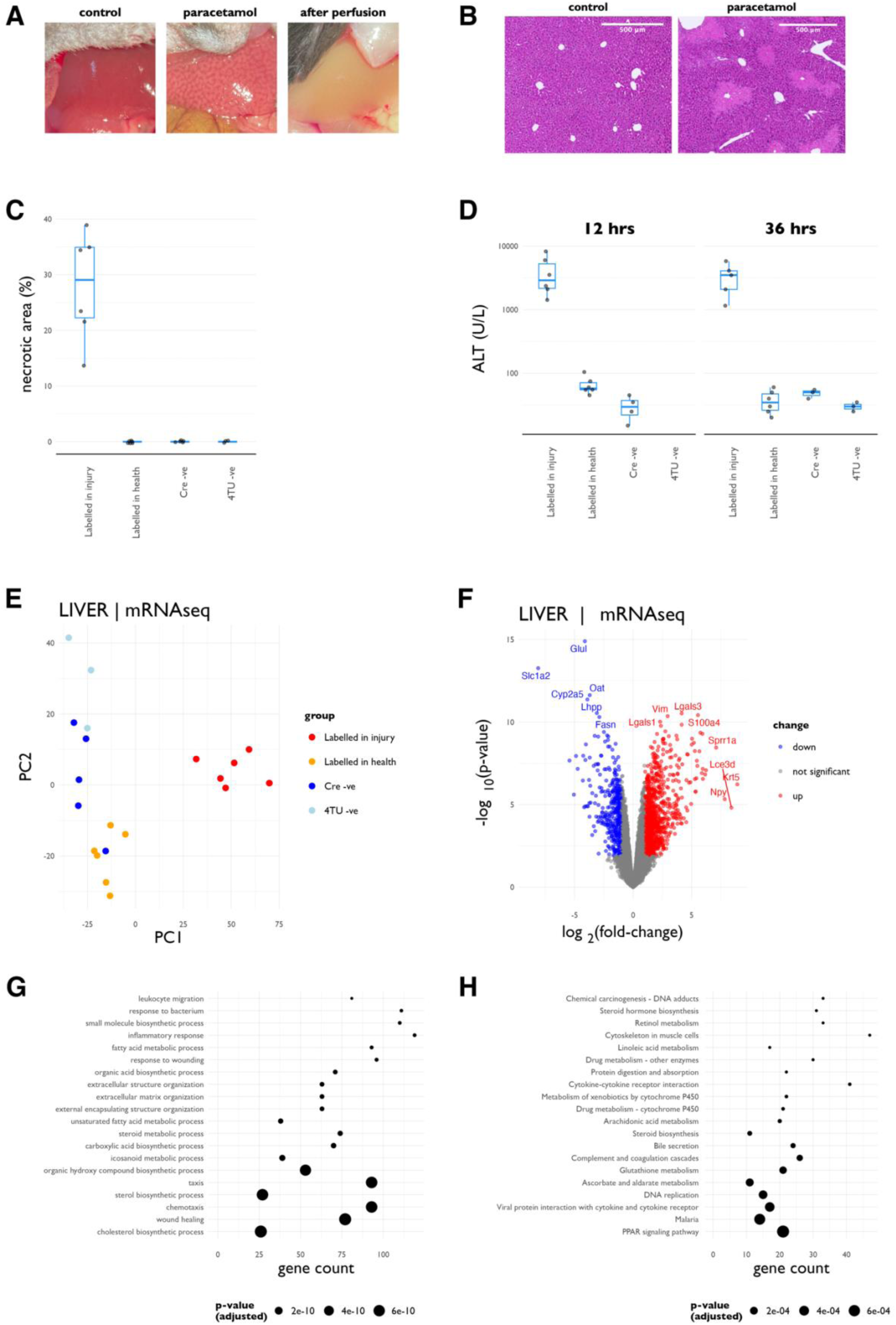
Paracetamol overdose induces acute hepatocellular injury. **A)** *Macroscopic pathology.* Representative photographs taken immediately before systemic perfusion. Necrosis was evident under the low-power dissecting microscope. **B)** *Liver histology.* H&E stain of fixed liver tissue showing necrosis after paracetamol administration. **C)** *Necrotic area.* Quantification of the necrotic area observed on H&E sections. **D)** *ALT*. Plasma ALT concentration at 12- and 36-hours post-paracetamol. **E)** *PCA.* Principle component analysis of the RNAseq data after liver injury. There were distinct global changes in the transcriptome of the liver injury group. **F)** *Volcano plot.* Differentially regulated genes in the liver after paracetamol injury. Genes that were differentially regulated with a fold-change exceeding 2 and a false-discovery rate of < 0.05 are depicted in red (if upregulated in injury) or blue (if downregulated). **G)** *GO analysis.* Pathways upregulated in liver after paracetamol injury, determined by gene ontology analysis. **H)** *KEGG analysis.* Pathways upregulated in liver after paracetamol injury in a KEGG analysis. Data from male mice; n = 6 (labelled after paracetamol), n = 6 (labelled in health), n = 5 (Cre-negative control), n = 3 (4TU-negative control).

The RNA labelling protocol achieved robust labelling of liver mRNA (Fig 5A&B) and small RNA (Fig 5E-I) in the context of paracetamol overdose. T>C conversion rates were, again, markedly lower in the small RNA SLAMseq dataset than in the mRNA dataset (Table 1). In a gene-wise analysis, 92% of all mRNAs and 59% of small RNAs within the liver were labelled in the context of acute liver injury, compared to 89% of mRNAs and 57% of small RNAs labelled using the same labelling protocol under healthy conditions (Table 2; supplemental Tables S8-9). There was universal labelling of known hepatocyte genes under healthy conditions (Fig 5C). However, labelling was not restricted to known hepatocyte “marker” genes (Fig 5C) and labelling in non-hepatocyte marker genes was more common after liver injury (Fig 5D). The known hepatocyte-enriched miRNA, miR-122, was labelled more strongly after paracetamol treatment, indicating increased rates of synthesis within surviving hepatocytes; there was no increase in labelling of known endothelial-enriched miRNAs (Fig 5J).

**Fig 5.**
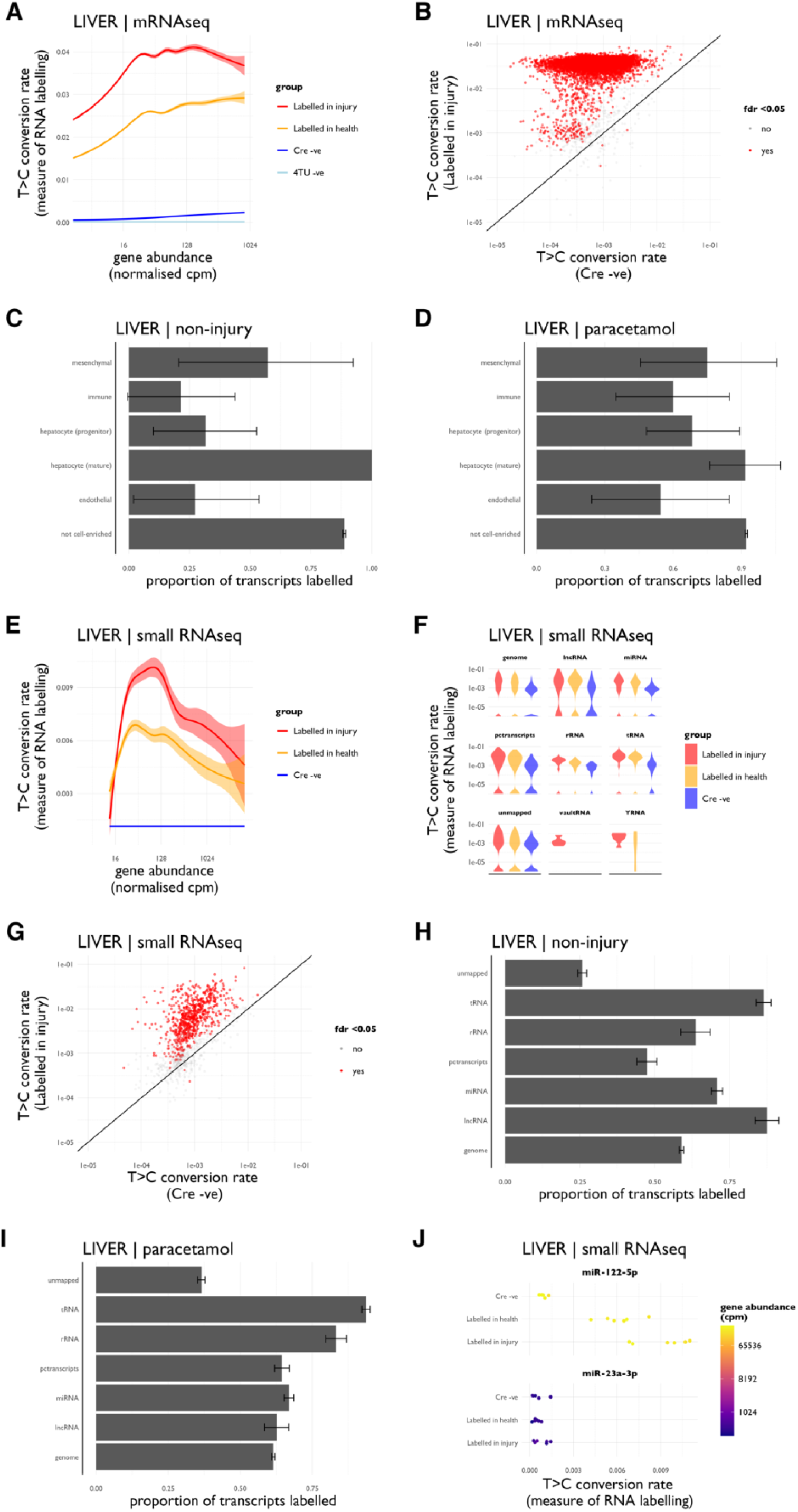
RNA labelling in the liver after injury. **A)** *T>C conversion rates.* T>C conversion rates in mRNA SLAMseq data. **B)** *Labelled transcripts.* Significantly labelled transcripts after liver injury in mRNA SLAMseq data. **C)** *Labelling of known marker genes.* Labelling rates within groups of “marker genes”, known to be enriched in defined cell types, as determined in liver mRNAseq data in health. **D)** *Labelling of marker genes in injury.* Labelling of marker genes in liver mRNAseq data after paracetamol. **E)** *T>C conversion rates.* T>C conversion rates in smallRNA. **F)** *T>C conversion rates.* T>C conversion rates in small RNA, stratified by RNA biotype. **G)** *Labelled transcripts.* Significantly labelled transcripts after liver injury in small RNA. **H)** *Labelling of small RNA in healthy liver.* Stratified by RNA biotype. **I)** *Labelling of small RNA in the liver after paracetamol.* Stratified by RNA biotype. **J)** *Labelling of miR-122 and miR-23a.* T>C conversion rates in the known hepatocyte-enriched miRNA, miR-122, and the known endothelial-enriched miRNA, miR-23a. Data from male mice; n = 6 (labelled after paracetamol), n = 6 (labelled in health), n = 5 (Cre-negative control), n = 3 (4TU-negative control).

**Table 1.** T>C conversion rates in liver injury experiment. ND = not determined. * p = 0.05; ** p < 10^−12^ for comparison to the Cre-negative group within that tissue by Wilcoxon signed rank test after Kruskal-Wallis rank sum test.

| tissue | group | mRNA |  |  | small RNA |  |  |
| --- | --- | --- | --- | --- | --- | --- | --- |
|  |  | *not labelled* | *labelled* | % | *not labelled* | *labelled* | % |
| **liver** | *labelled in health* | 1,365 | 10,555 | 88.6 | 353 | 462 | 56.7 |
| **liver** | *labelled in injury* | 976 | 11,205 | 92.0 | 520 | 753 | 59.2 |
| **liver** | *4sU* | 11,246 | 1,059 | 8.6 | ND | ND | ND |
| **kidney** | *labelled in health* | 12,701 | 618 | 4.6 | 1544 | 0 | 0.0 |
| **kidney** | *labelled in injury* | 8,794 | 4,535 | 34.0 | 1454 | 0 | 0.0 |
| **kidney** | *4sU* | 13,225 | 473 | 3.5 | ND | ND | ND |

**Table 2.** RNA labelling rates in the liver injury experiment. The number of mRNAs and small RNAs (i.e. distinct genes) labelled in liver and kidney in different experimental groups. ND = not determined. Labelling frequency of mRNA was significantly different between all experimental groups in both liver and kidney (p < 10^− 7^ by Chi-squared test for all between-group comparisons). Labelling frequency of small RNA was not different between experimental groups in the liver (p = 0.28) or the kidney (no labelling in either group).

| group | T>C conversion % (median, IQR) |  |  |  |
| --- | --- | --- | --- | --- |
|  | mRNA |  | small RNA |  |
|  | *liver* | *kidney* | *liver* | *kidney* |
| *4TU-negative* | 0.000 (0.000 - 0.015) * | 0.000 (0.000 - 0.023) ** | ND | ND |
| *Cre-negative* | 0.017 (0.000 - 0.116) | 0.000 (0.000 - 0.036) | 0.056 (0.013 - 0.120) | 0.064 (0.019 - 0.118) |
| *labelled in health* | 1.861 (0.595 - 2.821) ** | 0.027 (0.000 - 0.065) ** | 0.210 (0.049 - 0.705) | 0.051 (0.000 - 0.099) ** |
| *labelled in injury* | 3.304 (0.938 - 4.630) ** | 0.053 (0.000 - 0.105) ** | 0.300 (0.062 - 0.990) | 0.046 (0.000 - 0.093) ** |
| *4sU* | 0.000 (0.000 - 0.036) ** | 0.003 (0.000 - 0.034) ** | ND | ND |

### Labelled RNA is detected in the kidney after acute hepatocellular injury

After liver injury, we detected increased labelled mRNA in the kidney. T>C conversion rates were significantly higher than those observed under healthy conditions (median 0.053 *vs.* 0.027% T>C conversion; p < 10^−12^; Table 1; Fig 6A&B). Labelling was observed in known hepatocyte marker genes, to a greater extent than for marker genes of other liver cell types (Fig 6C&D). In contrast, there was no labelled small RNA within the kidney (Fig 6E-F; Table 2). The liver-specific miRNA, miR-122, did not exhibit increased rates of T>C conversion within kidney tissue (Fig 6G).

**Fig 6.**
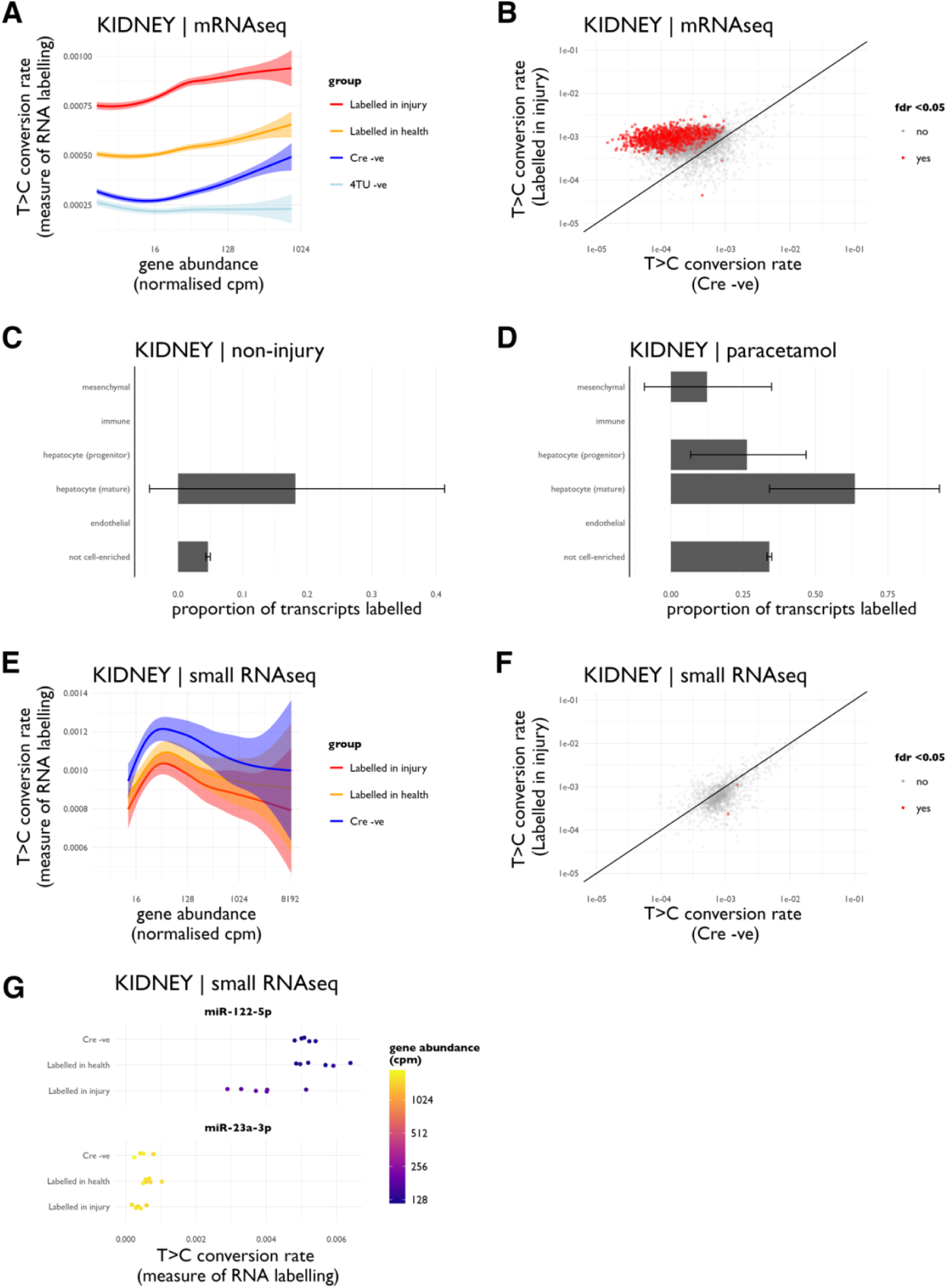
Labelled mRNA in the kidney after liver injury. **A)** *T>C conversion rates in kidney.* T>C conversion rates in kidney RNA after liver injury, in the SLAMseq mRNA data. **B)** *Labelled transcripts in kidney.* Labelled transcripts in kidney mRNA SLAMseq data after liver injury. Labelled mRNA transcripts were detected by comparing gene-wise T>C conversion rates between Cre-positive (RNA labelling) and Cre-negative (control) groups using the beta-binomial method and setting a false discovery rate of 0.05. **C)** *Labelling of liver marker genes.* Labelling rates within groups of “marker genes”, known to be enriched in defined liver cell types, as determined in kidney mRNAseq data in health. **D)** *Labelling of liver marker genes in injury.* Labelling of marker genes in kidney mRNAseq data after paracetamol. **E)** *T>C conversion rates in kidney.* T>C conversion rates in kidney small RNA after liver injury. **F)** *Labelled transcripts in kidney.* There were no labelled transcripts in kidney small RNA after liver injury. **G)** *Labelling of miR-122 and miR-23a.* T>C conversion rates in the known hepatocyte-enriched miRNA, miR-122, and the known endothelial-enriched miRNA, miR-23a. Data from male mice; n = 6 (labelled after paracetamol), n = 6 (labelled in health), n = 5 (Cre-negative control), n = 3 (4TU-negative control).

### Characteristics of labelled mRNA in the kidney

Our findings suggest that thiolated RNA may be transferred from liver to kidney, either as intact, potentially functional RNA molecules or as dissociated nucleotides that are subsequently incorporated into nascent mRNA in kidney cells. Although SLAMseq cannot directly distinguish between these possibilities, we can draw inferences about which is more likely by examining which kidney RNAs are labelled under different conditions. We reasoned that indiscriminate labelling of kidney RNA by dissociated thiolated nucleotides would preferentially label RNA molecules transcribed at higher rates within kidney cells, hence targeting transcripts that are more abundant in kidney. Conversely, transfer of intact RNA molecules from hepatocytes would result in 4TU-dependent labelling within genes known to be more abundant within hepatocytes. However, this analysis is confounded by a strong positive correlation between abundance in liver and kidney for most genes (Fig 7E). We therefore included a group of mice treated with 4-thiouridine (4sU). Whereas 4TU is expected to be incorporated into nascent RNA only in UPRT-expressing hepatocytes, 4sU will be incorporated into nascent RNA in all cells (supplemental Fig S1C). Therefore, the 4sU-treated group provides important context, enabling comparison to a dataset in which kidney transcripts are *known* to be labelled by dissociated nucleotides.

**Fig 7.**
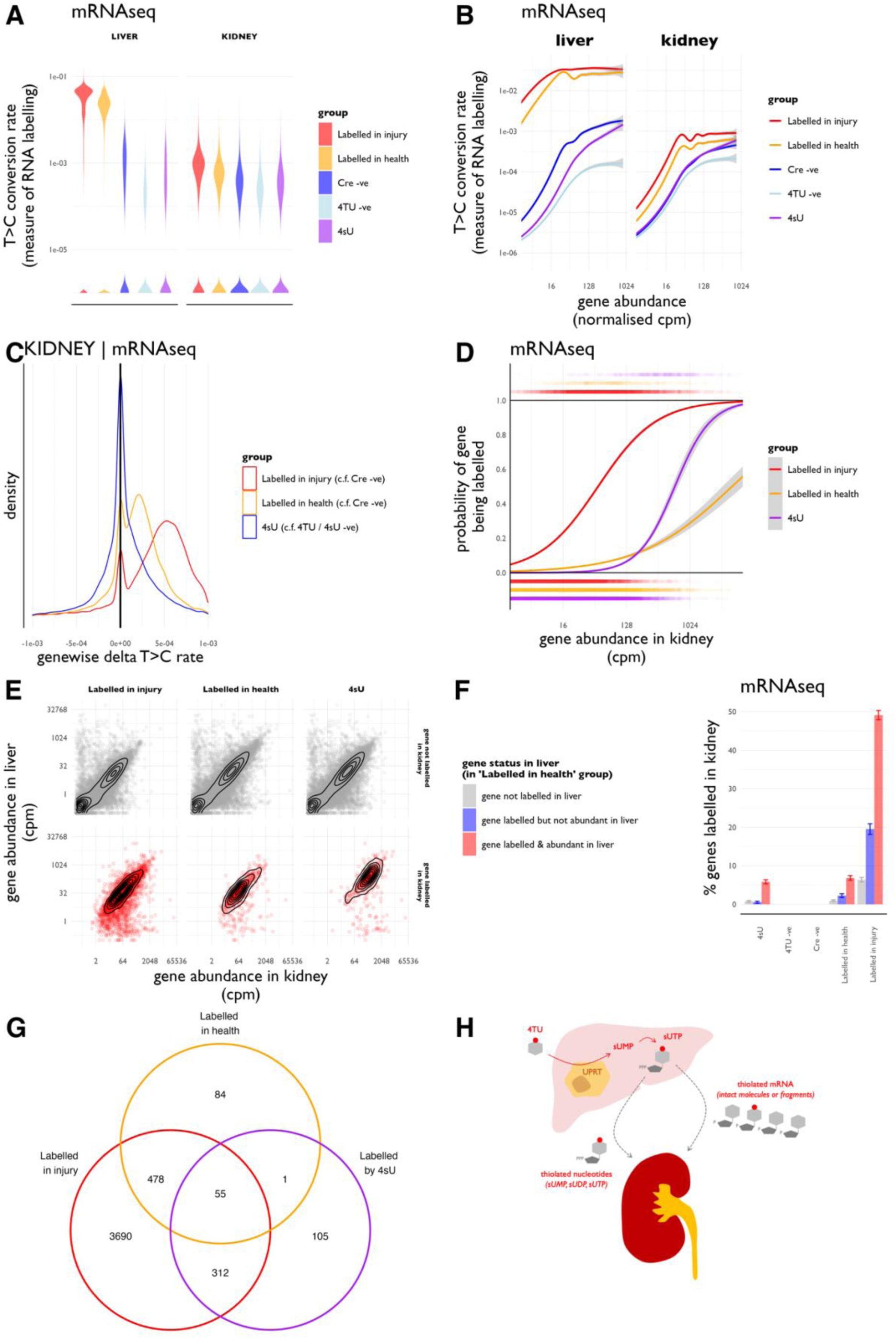
Characterisation of labelled kidney transcripts. **A)** *Indiscriminate labelling with 4-thiouridine.* Mice were administered with 4-thiouridine (4sU) to induce labelling of all nascent RNA in all cell types. The dose used induced a low level of T>C conversion in both liver and kidney, approximately equivalent to that seen in the Cre-negative, 4TU-exposed control group. **B**) *T>C conversion rates in the universal labelling group, dependence on transcript abundance.* Data are the same as for Fig 5A & Fig 6A, but the 4sU group has been added and the results are plotted on a log scale. **C**) *Gene-wise difference in T>C conversion rates.* The difference in T>C conversion rate between labelling and control groups was determined for each gene. The relevant control group was the Cre-negative, 4TU-exposed group for the “labelled in injury” and “labelled in health” groups and the 4sU, 4TU-naïve group for the “4sU” group. **D)** *Probability of any given gene being labelled within the kidney mRNA SLAMseq dataset.* Each point below zero represents a gene that is unlabelled; each point above 1 represents a labelled gene. The probability of any gene being labelled within each experimental group was determined by fitting a sigmoidal curve. **E)** *Relationship between gene abundance in liver and kidney, stratified by labelling status within kidney*. **F**) *Proportion of genes labelled in the kidney, stratified by gene status in liver.* Genes were classified according to their labelling status in the liver in the “labelled in health” group. A gene was classified as being labelled in the liver according to the beta-binomial test depicted in Fig 5B it was deemed abundant in liver if it was detected with an adjusted cpm > 10. The proportion of genes identified as being labelled within the kidney was then determined, stratifying the results according to whether the gene was also labelled in liver. **F**) *Venn diagram showing number of genes labelled in kidney in different experimental groups.* **G**) *Concept figure showing model compatible with our observations*. Our data are compatible in a model in which there is transfer of dissociated thiolated nucleotides and intact RNA molecules or large fragments of RNA molecules from liver to kidney, which is increased after acute hepatocellular injury. Data from male mice; n = 6 (labelled after paracetamol), n = 6 (labelled in health), n = 5 (Cre-negative control), n = 3 (4TU-negative control), n = 3 (4sU positive control).

4sU administration induced a low level of RNA labelling, approximately equivalent to that seen in the Cre-negative, 4TU-exposed group (Fig 7A&B). The distribution of T>C conversions in the kidney differed between the 4sU-treated and 4TU-treated groups; 4sU treatment induced a smooth right-skew in genewise delta T>C values (i.e. T>C conversion rate in 4sU-treated minus 4sU-negative groups) whereas 4TU treatment induced a bimodal distribution, in which a distinct pool of transcripts were clearly labelled (Fig 7C).

The probability of any kidney gene being labelled increased with gene abundance within kidney tissue. This was true for all experimental groups, but the shape of this relationship differed between groups (Fig 7D). In genes that were present at lower abundance within kidney, the probability of being labelled by 4TU in health was greater than the probability of being labelled by 4sU, despite this dose of 4sU inducing much higher labelling rates in genes at higher abundance. This observation is compatible with the transfer of intact RNA molecules from liver to kidney, detected as labelling in a subset of low-abundance transcripts in kidney tissue.

Gene abundance within kidney and liver tissue was strongly correlated for most genes (Fig 7E). The shape of this relationship was broadly similar within the sets of unlabelled and labelled kidney genes (Fig 7E). Those transcripts that were labelled and relatively abundant (cpm > 10) in uninjured liver were more likely to be labelled within the kidney in the 4TU-treated groups. They were also most likely to be labelled in the 4sU-treated group, likely reflecting the fact that these transcripts are also abundant – and hence highly transcribed – within the kidney. However, transcripts that were labelled but not abundant (cpm < 10) in uninjured liver were more likely to be labelled within the kidney in 4TU-treated mice than were transcripts that were not labelled in liver; this was not the case in 4sU-treated mice (Fig 7F). This difference between 4sU- and 4TU-treated mice is compatible with transfer of intact RNA molecules from liver to kidney in 4TU-treated mice.

### Predicted function of RNA transferred from liver to kidney

618 transcripts (4.6%) were labelled in the kidney after treatment with 4TU under healthy conditions. This increased to 4535 transcripts (34.0%) in the context of acute hepatocellular injury. In the 4sU-treated group, 473 transcripts (3.5%) were labelled in the kidney (Table 2; Fig 7G). Those transcripts labelled within the kidney in the 4TU and 4sU-treated groups are listed in supplemental Table S10.

Using pathway analyses, we examined the known functions of mRNAs that were likely transferred from liver to kidney. We reasoned that the set of 478 mRNAs that were labelled by 4TU in the kidney under both basal and injury conditions, but not by 4sU, was likely to be enriched in mRNAs that are transferred from liver to kidney. In gene ontology and KEGG analyses, these RNAs were enriched for terms relating to RNA transcription and processing, as well as other diverse biological pathways (supplemental Fig S3 A&B). Similarly, we reasoned that the set of 3690 mRNAs that were labelled in kidney only in 4TU and paracetamol-treated mice, was likely to be enriched in mRNAs that are transferred from liver to kidney after liver injury. Within this set, there was over-representation of mRNAs associated with catabolic and injury processes such as autophagy, mitophagy and ferroptosis (supplemental Fig S3 C&D).

## Discussion

### RNA is transferred from liver to kidney

Our main finding is that RNA is transferred *en masse* from the liver to the kidneys in mice. The extent to which exRNA signalling serves a physiological purpose is controversial,^6^ with evidence largely derived from experiments in which single miRNAs are perturbed^13,14,19–23^. Here, we provide evidence of one necessary pre-condition of exRNA signalling: the transfer of RNA between cells. Our results suggest that any intercellular signalling would be mediated by multiple RNAs, rather than a single dominant miRNA. Whether such transfer is sufficient to induce meaningful biological effects in the recipient tissue (in this case the kidney) under physiological conditions, remains an important open question.

### A SLAMseq approach was able to track mobile extracellular RNAs

We sought to label RNA in hepatocytes with 4TU and then detect labelled RNA in liver and kidney using SLAMseq. The success of this approach is critically dependent on stringent hepatocyte-specific expression of Cre recombinase (and therefore of uracil phosphoribosyltransferase). The AAV8-TBG-Cre vector, through the combined effects of viral tropism and the Thyroxine Binding Globulin promoter, has been repeatedly shown to induce hepatocyte-specific Cre expression.^24^ We confirmed that there was no off-target expression of Cre and UPRT in the kidney at the level of genomic DNA, RNA and protein. We demonstrated that RNA labelling (marked by T>C conversions) was evident in liver RNA, specifically labelling known hepatocyte marker genes as well as most other (non-cell type-restricted) transcripts. We were also able to detect T>C conversions in kidney RNA.

SLAMseq has been used previously in an attempt to tag and track small non-coding RNAs moving from the epididymis to developing sperm.^25^ Although this prior attempt provided an elegant proof-of-principle that SLAMseq can be used to track exRNA moving between tissues, analysis was restricted to miRNAs in two biological replicates and lacked the resolution to identify *which* miRNAs were being transferred. Moreover, that study was not able to differentiate between the intercellular transfer of intact RNA molecules and dissociated thiolated nucleotides. By using a greater number of biological replicates and by comparing which set of RNAs were labelled in different states (health, liver injury, universal labelling with 4sU), we were able to identify those RNAs that were transferred from liver to kidney with moderate confidence.

### RNA is likely to be transferred intact and as dissociated nucleotides

After using 4TU to label RNA specifically in hepatocytes, we detected labelled (thiolated) RNA in the kidney. This observation might result from any of three phenomena:

i. the off-target expression of UPRT in the kidney;
ii. the transfer of dissociated ribonucleotides (4T-UMP, 4T-UDP, 4T-UTP) from liver to kidney;
iii. the transfer of intact thiolated RNA molecules.

(There could also be transfer of intact RNA that is then degraded within kidney cells, and the nucleotides re-cycled into nascent RNA molecules; our approach is unable to distinguish between this and transfer of dissociated nucleotides.)

In addition to our data showing no off-target expression of Cre and UPRT, our finding of increased rates of labelling in kidney RNA after acute hepatocellular injury is strongly suggestive of transfer of RNA from liver to kidney. Our results suggest that there is transfer of both intact RNA molecules (or large molecular fragments) and dissociated nucleotides from liver to kidney.

Compatible with dissociated nucleotide transfer, RNA labelling in the kidney was not restricted to RNAs that were labelled in liver. Indeed, there was near-universal labelling of high-abundance kidney mRNAs after liver injury, suggesting that the transfer of dissociated nucleotides exerts a strong effect on kidney RNA labelling in this context.

Several lines of evidence suggest transfer of large RNA molecules from liver to kidney. First, labelling was evident in kidney transcripts mapping to known hepatocyte “marker genes”, more than for marker genes of other liver cell types. Second, a distinct population of relatively low-abundance labelled transcripts was detected in healthy mice exposed to 4TU but not in mice treated with 4sU – despite using a dose of 4sU that induced higher labelling rates in high-abundance transcripts. Third, those transcripts that were labelled but not abundant in liver were more likely to be labelled in the kidneys of 4TU-treated mice, but not so in 4sU-treated mice – despite similar labelling rates of genes that are not labelled in liver in those experimental groups.

Consistent with both intact RNA and dissociated nucleotides being transferred, we observed a set of transcripts that were labelled in the kidney by both 4TU and 4sU treatments as well as distinct sets that were labelled only by 4TU.

### Labelled mRNA was detected in kidney but labelled small RNA was not

exRNA researchers have largely studied small non-coding RNAs because these are enriched in extracellular vesicles (EVs) and other exRNA carriers.^26^ However, mRNAs are consistently detected within exRNA preparations and have been shown to move between cells *in vivo* and to exert biological effects in target cells.^27–31^ Some extracellular mRNAs have been observed to enter the nucleus of recipient cells.^32^ mRNAs appear to be selective packaged into EVs, in part through a mechanism dependent on their binding to the heterogenous nuclear ribonucleoprotein, HNRNPA2B1.^33^

The mRNAs that we identified as being likely transferred from liver to kidney were enriched for GO terms relating to RNA transcription and processing. Intriguingly, this same enrichment for RNA-processing-related GO terms has been observed in EV-associated mRNAs.^33^ Therefore, our findings extend observations that have been made by multiple investigators in diverse contexts, and suggest that exRNA signalling – if it operates in mammals – is unlikely to be restricted to small non-coding RNAs.

Perhaps surprisingly, we did not detect labelled small RNA within the kidney. This might reflect a lack of small RNA transfer from liver to kidney or that our approach was not sufficiently sensitive to detect this. The latter is plausible. In the liver, our labelling protocol induced mean rates of T>C conversion that were ∼ 5-fold lower for small RNAs than for mRNA. Labelling rates for small RNA were comparable to those observed elsewhere in the literature (e.g. 0.15 % was observed by Sharma et al. after labelling small RNA in epididymis^25^). This relatively lower rate of small RNA labelling likely reflects the increased stability of small RNAs compared to mRNA.^34,35^ This less extensive labelling of liver small RNA, coupled with the lower number of T residues typically present per ∼22nt miRNA than in the 3’UTR of a typical mRNA, will have made it much harder for our SLAMseq approach to confidently identify labelled miRNA (compared to mRNA) in kidney tissue.

### Perspectives

Our results provoke further questions and hypotheses concerning the physiological consequences of liver-to-kidney exRNA signalling. After liver injury, labelled RNA in the kidney was enriched for mRNAs participating in cellular injury pathways such as autophagy, mitophagy and ferroptosis. This is consistent with the hypothesis that hepatocyte injury provokes the release of RNAs participating in “injury-response” pathways that are then taken up by kidney cells, activating the same injury response pathways in the kidney. Such a phenomenon might be evolutionarily advantageous in the context of a pervasive acute injury (e.g. exposure to a toxin) but could be maladaptive / pro-fibrotic in response to chronic injury. It would be productive to determine to extent to which these transferred RNAs induce physiologically relevant phenotypic changes in kidney cells, and whether these could plausibly mediate the well-established association between chronic liver and chronic kidney disease.^17^

### Conclusion

Using SLAMseq, we have demonstrated transfer of thiolated ribonucleotides from liver to kidney in the mouse and shown that this transfer is augmented by acute hepatocellular injury. We conclude that there is transfer of both dissociated nucleotides and intact, potentially functional, mRNA molecules.

## Methods

### Transgenic mice

All procedures were performed under UK Home Office licence in accordance with the Animals (Scientific Procedures) Act, 1986 and after review and approval by local veterinary surgeons. Mice had free access to water and a standard RM1 diet (Special Diet Services, Witham, UK) and were maintained on a 12 h–12 h light–dark cycle (lights on at 7 am).

Floxed-stop-UPRT mice^36^, B6;D2-Tg(CAG-GFP,-Uprt)985Cdoe/J were purchased from Jackson laboratories and were maintained through homozygous-homozygous crosses. Mice were genotyped by Transnetyx, using PCR primers that we designed to amplify products across the 3’ end of the transgene locus on chromosome 12.^37^ Podocin-UPRT mice were generated by crossing homozygous floxed-stop-UPRT females and hemizygous podocin-Cre males (Tg(Nphs2-cre)1Nagy).^38^

### AAV8-TBG-Cre treatment

The AAV8.TBG.PI.Cre.rBG vector (Addgene #107787) was diluted to 6.25 × 10^12^ genome copies per microlitre in sterile PBS and stored in small aliquots at -70 °C. Immediately after thawing, it was diluted 10-fold in sterile PBS and 100 mcl was injected *via* the lateral tail vein, delivering a dose of 6.25 × 10^10^ genome copies per mouse. Mice were left for at least four weeks to allow efficient expression of recombinant UPRT before 4-thiouracil administration.

### Genomic DNA recombination assay

Tissue samples were digested with proteinase K (500 mcg/ml in 60 mM Tris, 100 mM EDTA, 0.5% SDS at 55°C for 16 hrs) and then genomic DNA was isolated by phenol-chloroform extraction and isopropanol precipitation, including an RNAse step (25 mcg/ml RNAseA for 30 mins at 37°C). 40 ng (c. 1.4 × 10^4^ genome copies) gDNA was used as a template in PCR reactions designed to amplify products from the intact or recombined floxed-stop-UPRT locus (supplemental FigS2A). A common forward primer (CGTGCTGGTTATTGTGCTGT) was used with two reverse primers designed respectively to amplify a 332 bp region from the intact locus (AAGTCGTGCTGCTTCATGTG) or a 167 bp from the recombined locus (TCTCGACAAGCCCAGTTTCT). The latter primer could also amplify a 1707 bp region from the intact locus and therefore we used a relatively fast cycling program: 24 cycles of 94 °C × 45s, 58 °C × 15s, 72 °C × 15s, after a 12-cycle touch-down phase in which the annealing temperature was reduced by 1°C per cycle until landing on 58 °C.

### 4-thiouracil and 4-thiouridine treatment

4-thiouracil (4TU, Sigma 440736) was dissolved at 200 mg/ml (1560 mM) in DMSO and stored in small aliquots at –20°C. Immediately prior to administration, this was diluted 1:10 in corn oil to give a 20 mg/ml (156 mM) solution in 90% corn oil / 10% DMSO. Mice received three 20 ml/kg body weight (= 400 mg/kg = 3.12 mmoles per kg) injections either subcutaneously or intraperitoneally according to the protocols in supplemental FigS1.

4-thiouridine (4sU, Sigma T4509) was dissolved at 40.6 mg/ml (156 mM) in DMSO and stored in small aliquots at –20°C. Immediately prior to administration, this was diluted 1:10 in 0.9% NaCl to give a 15.6 mM solution in 10% DMSO. Mice received three 20 ml/kg (= 81 mg/kg = 0.312 mmoles per kg) injections subcutaneously, at 33, 24 and 12 hours prior to cull.

### Paracetamol-induced liver injury

Mice were fasted for 12 hrs from 9 am prior to paracetamol administration. Paracetamol (Sigma) was dissolved in hot, sterile 0.9% NaCl at 15 mg/ml and delivered by IP injection (20 ml/kg body weight = 300 mg/kg). Thereafter, mice were housed in an incubator (27°C, 40% humidity) and provided with supplementary mashed diet, to minimise paracetamol-induced morbidity.

### Blood and tissue sampling

Blood samples (50 – 100 mcl) were obtained at 12 hrs after paracetamol injection by lateral tail-vein sampling. At the end of the study, mice were euthanised by terminal perfusion under anaesthesia (pentobarbital sodium 75 – 150 mg per kg body weight by IP injection).

Through a mid-line laparotomy, a segment of aorta was clamped inferior to the origin of the renal arteries. The aorta was then cannulated with “P10” BTPE tubing (outer diameter 0.61 mm), secured with a silk suture. The clamps were removed, and a terminal blood sample taken before the inferior vena cava was vented. Ice-cold PBS (30 – 50 ml) was perfused through the aorta until the exit fluid ran clear and all organs were visibly well-perfused (Fig 4A).

Organs were harvested in strict order (kidneys then spleen then heart then liver), using single-use forceps and scalpels so that there was no cross-contamination from liver to other organs. Tissue samples for RNA extraction were incubated in RNALater for 18 hrs at 4°C before being stored at -70°C; tissue samples for DNA and protein extraction were snap-frozen and stored at -70°C. Tissue samples for histology were fixed in 10% neutral buffered formalin (= 4% formaldehyde) for 24 hrs at 4°C before being embedded in paraffin, sectioned and stained with H&E.

Blood samples were collected into K-EDTA tubes and centrifuged for 2000 *g* for 10 mins before storing plasma at -70 °C. Plasma alanine transaminase (ALT) activity was measured in a kinetic assay,^39^ using a commercial kit (Sentinel Diagnostics 17234H) adapted for use on either a Cobas Fara or Mira analyser (Roche Diagnostics). Intra-assay and inter-assay precision were CV < 4% and CV < 8% respectively.

### RNA extraction and alkylation

RNA was extracted and alkylated as per published SLAMseq protocols.^40,41^ Briefly, tissue samples were lysed in a monophasic phenol / guanidine isothiocyanate solution (TRIsure) and then RNA extracted by chloroform extraction and isopropanol precipitation in the presence of 20 mcg glycogen and 0.1 mM DTT. Samples were treated with DNAse (Thermo AM1907) and then cleaned on silica columns (Zymo R1013), eluting into 15 microL 1 mM DTT. RNA samples were alkylated by treating with 10 mM iodoacetamide in 50 mM sodium phosphate buffer / 50 % DMSO for 50 °C for 15 mins. The reaction was quenched with 20 mM DTT and then RNA samples purified by ethanol precipitation. The integrity and concentration of alkylated RNA samples was assessed by automated capillary electrophoresis (on a LabChip GX Touch). To verify success of the alkylation reaction, 1 mM 4-thiouracil controls were included and diluted 1:10 before being subjected to UV absorbance spectrophotometry (Nanodrop in UV-Vis mode).

### RNA sequencing and quantification of T>C conversions

Alkylated RNA samples were submitted to Lexogen who prepared and sequenced libraries. RNAseq data were deposited in the European Nucleotide Archive (ENA) at EMBL-EBI under accession number PRJEB75334 (https://www.ebi.ac.uk/ena/browser/view/PRJEB75334; embargoed until paper publication). Read counts are listed in supplemental Table S13.

For mRNAseq, 500 ng input total RNA was amplified in 13 cycles of PCR during library preparation with a QuantSeq-FWD kit v2. (Quantseq is a method for sequencing the 3’ ends of mRNA.^42^) Sequencing was performed on an Illumina NextSeq 2000 sequencer, to generate single-end 100 bp reads. To map and quantify T>C conversions in the 3’UTRs of mRNA, RNAseq data were analysed using the SLAMDUNK analysis pipeline.^40,43^

For smallRNAseq, 200 ng input total RNA was amplified in 18 – 24 cycles of PCR during library preparation with a Perkin Elmer NextFlex Small RNA-Seq kit v3. Sequencing was performed on an Illumina NextSeq 2000 sequencer, to generate single-end 100 bp reads. To map and quantify T>C conversions in small RNAs, we used our “smallSLAM” analysis pipeline (https://github.com/robertwhunter/smallSLAM; https://zenodo.org/doi/10.5281/zenodo.11083131). This accounts for the high number of duplicated reads and unique mapping requirements of smallRNA sequencing data. Reads were trimmed using cutadapt, retaining reads with a length exceeding 15 bases. Unique reads were then grouped into families comprising one “parent” sequence and multiple “child” sequences containing one or more T>C conversions. The T>C conversion rate was computed for each parent sequence. Reads were mapped sequentially to rRNA, tRNA, miRNA, piRNA, snoRNA, snRNA, vaultRNA, YRNA, lncRNA, protein-coding RNA and whole genome reference sequences (supplemental Table S12). Mapping was performed using Bowtie2, permitting 1 mismatch in an 18-base seed.

### Analysis of RNA labelling and differential expression

Labelled RNAs were identified by making gene-wise, between-group comparisons of T>C conversion rates using a beta binomial test (countdata package version 1.3).^44^ To define a set of known cell-enriched (or “marker” genes) we mined the CellMarker database.^45^ We selected marker genes known to be expressed in normal (i.e. non-cancer) cell types in mouse liver (supplemental Table S11).

Differential expression analysis was performed using edgeR (v3.40.2).^46^ Low-abundance reads were filtered out and then counts were normalized using the TMM (trimmed mean of M-values) method. Differential expression was determined using a 2-group generalized linear model. Transcripts were deemed to be differentially expressed if their expression was at least 2-fold different between groups and false-discovery rate was less than 0.05. Gene ontology and KEGG pathway analysis was performed using the clusterProfiler package (version 4.6.2).^47^

### RNA biotinylation and dotblot

10 mcg total RNA (extracted without exposure to DTT) was biotinylated by mixing with 1 mcg MTSEA-biotin-XX (Biotium) in 10 mM HEPES, 1 mM EDTA, 20% DMF, pH 7.5 for 120 minutes at room temperature. RNA was then cleaned by chloroform extraction in phase-lock gel tubes (MaxTRACT high density tubes, Qiagen) and ethanol precipitation to remove unbound biotin. Biotinylated RNA was heated to 65°C for 5 minutes and then cooled on ice, before being spotted in 2.5 mcl drops onto a dry nylon membrane (Hybond-N+, Roche 11417240001). Biotinylated oligo-dT (Promega Z5261) was used as a positive control. The membrane was dried for 30 minutes then wet in 10x SSC (1.5M NaCl, 150 mM tri-sodium citrate) before the RNA was cross-linked by UV light (1200 mJ cm^−2^ on Spectrolinker XL-1500). Membranes were blocked for 30 minutes in 125 mM NaCl, 17 mM Na_2_HPO_4_, 7.3 mM NaH_2_PO_4_, 1% SDS and then incubated with Streptavidin-HRP (Amersham RPN1231, 1:2000 in blocking buffer) for 10 minutes. Membranes were then washed in a 1:10 dilution of blocking buffer (twice for 20 minutes) then in 100 mM Tris, 100 mM NaCl, 21 mM MgCl_2_, pH 9.5 (twice for 5 minutes). HRP signal was detected using ECL reagent and a LICOR Odyssey Fc imager.

### Immunoblot

Quarter kidneys, containing cortex and medulla, were homogenised in 250 mM sucrose / 20 mM triethanolamine with protease inhibitors (1% Merck Protease Inhibitor Cocktail III). The homogenate was cleared of large debris by centrifugation at 4000 *g* for 15 mins at 4 °C. Samples were prepared for SDS-PAGE by mixing with 2x tris-glycine SDS sample buffer (Invitrogen) and DTT (final concentration 50 mM) and then heating to 70 °C for 15 mins. SDS-PAGE was carried out using Novex WedgeWell 4-12% Tris-glycine gels. Gels were blotted onto PVDF by wet transfer in Tris-glycine buffer with 20% methanol. Membranes were then washed in 0.2% Tween in PBS and blocked in 5% (w/v) milk powder before being incubated with primary antibody (anti-HA, CST C29F4 or anti-βactin, CST 13E5 at 1:1,200) at 4 °C overnight. After three washes in 0.2% Tween in PBS, membranes were incubated with HRP-conjugated secondary antibody (mouse anti-rabbit IgG-HRP, sc-2357 at 1:10,000) for 60 mins at room temperature before being washed thrice again. HRP signal was detected using ECL reagent and a LICOR OdysseyM imager.

### Data analysis and statistics

Data were analysed in R (version 4.2.2)^48^ using the Tidyverse package (version 2.0.0)^49^.

To facilitate plotting of gene abundance on a log axis, we added a small “nudge” of 10^−1^ to all normalised counts per million (cpm), so that zero values could be plotted. Similarly, prior to plotting T>C conversion rates, we added a “nudge” of 10^− 6^.

We compared T>C conversion rates between groups using a Kruskal-Wallis rank sum test with *post hoc* two-sided, paired Wilcoxon signed rank tests with Bonferroni correction. We compared labelling frequencies between groups using a Pearson’s Chi-squared test with Yates’ continuity correction.

In adjusting for multiple comparisons in differential gene expression analysis or beta-binomial testing for labelled RNA, we deemed results to be statistically significant at a Benjamini-Hochberg false-discovery rate of 0.05. When determining the proportion of labelled genes within a given set, 95% confidence intervals were derived by the bootstrap method, sampling 10,000 times with replacement. We used the R boot package (v.1.3-28) and the “normal” method for constructing confidence intervals.^50^ If there were fewer than 10 genes in the set or the labelling rate within the set was 0 or 100%, we did not attempt to derive confidence intervals.

## Acknowledgements

This work was supported by grants from Wellcome (209562/Z/17/Z), BHF (RE/18/5/34216), Medical Research Scotland (ECG-1781-2022) and the China Scholarship Fund.

This work made use of the resources provided by the Edinburgh Compute and Data Facility (ECDF) (http://www.ecdf.ed.ac.uk/). We were assisted by the Edinburgh Bioquarter Shared University Research Facilities (SuRF) for ALT assays (Specialist Assay Service), RNA quantification (Biomolecular Core), histological processing and imaging (Histology Core).

We are grateful to Richard Coward (University of Bristol) for generously providing podocin-Cre mice and to Olivia Matthews for sharing images of AAV8-Tbg-Cre-injected mTmG mice. We are grateful to Nacho Vinuela-Fernandez, Kylie Matchett and Melisande Addison for advice in refining the paracetamol injury protocol. Sujai Kumar helped to design the smallSLAM pipeline for quantifying T>C conversions in small RNA SLAMseq datasets.

## Author contributions

**RWH** conducted most of the experimental work and analysis, wrote the first and final drafts of the paper and is the ultimate guarantor of data integrity. **JS, TP** conducted some of the experiments (gDNA recombination assay, HA-UPRT immunoblot, liver necrotic area quantification). **MAB, ND** and **AB** assisted in the design of SLAMseq experiments and data interpretation; they revised the manuscript in its final stages. **JWD** had critical oversight of experimental design and data analysis and revised the manuscript at early and final stages.

## Supplemental material

### List of supplemental tables

Table S1 – Group characteristics in small initial experiment.

Table S2 – Group characteristics in liver injury experiment.

Table S3 ** – T>C conversion rates in initial experiment.

Table S4 ** – RNA labelling rates in initial experiment.

Table S5 – mRNA labelled in the liver in initial experiment.

Table S6 – small RNA labelled in the liver in initial experiment.

Table S7 – mRNA labelled in the kidney in initial experiment.

Table S8 – mRNA labelled in the liver in liver injury experiment.

Table S9 – small RNA labelled in the liver in liver injury experiment.

Table S10 – mRNA labelled in the kidney in liver injury experiment.

Table S11 – Marker genes for cell types within the liver.

Table S12 – Reference genomes for small RNA mapping.

Table S13 ** – RNAseq dataset read counts.

Tables marked with ** are displayed below; the others are provided as standalone .csv files.

### List of supplemental figures

FigS1 – Experimental design.

FigS2 – Genomic DNA (gDNA) recombination assay.

FigS3 – Pathway analyses of mRNAs likely to be transferred from liver to kidney.

### Supplemental tables

**Table S3.** T>C conversion rates in initial experiment. * p = 0.0002; ** p < 10^−9^ for comparison to the Cre-negative group within that tissue by Wilcoxon signed rank test after Kruskal-Wallis rank sum test.

| group | T>C conversion rate (median, IQR) |  |  |  |
| --- | --- | --- | --- | --- |
|  | mRNA |  | small RNA |  |
|  | *liver* | *kidney* | *liver* | *kidney* |
| *RNA labelling* | 0.709 (0.043 - 4.474) ** | 0.009 (0.000 - 0.063) ** |  |  |
| *Cre-negative* | 0.000 (0.000 - 0.027) | 0.000 (0.000 - 0.032) |  |  |
| *UPRT-negative* | 0.000 (0.000 - 0.030) * | 0.000 (0.000 - 0.031) |  |  |
| *4TU-negative* | 0.000 (0.000 - 0.021) ** | 0.000 (0.000 - 0.030) ** |  |  |

**Table S4.** RNA labelling rates in initial experiment. The number of mRNAs and small RNAs (i.e. distinct genes) labelled in liver and kidney in the RNA labelling group.

| tissue | mRNA |  |  | small RNA |  |  |
| --- | --- | --- | --- | --- | --- | --- |
|  | *not labelled* | *labelled* | % | *not labelled* | *labelled* | % |
| **liver** | 10,451 | 1,501 | 12.6 | 1,291 | 327 | 20.2 |
| **kidney** | 13,500 | 81 | 0.6 | 1,265 | 0 | 0.0 |

**Table S13.** RNAseq read counts. The number of reads used to T>C quantification, after mapping, filtering and excluding SNPs, was smaller than the total number of reads in trimmed libraries.

| experiment | RNAseq modality | reads per trimmed library | reads used for T>C quantification |
| --- | --- | --- | --- |
| | | (mean ± SD) | (mean ± SD) |
| initial experiment in male and female mice | *mRNA (Quantseq)* | 21.7 ± 1.71 million | 10.1 ± 0.81 million |
| | *small RNA* | 22.5 ± 5.7 million | 18.5 ± 5.1 million |
| liver injury in male mice | *mRNA (Quantseq)* | 26.1 ± 3.68 million | 15.1 ± 2.02 million |
| | *small RNA* | 17.2 ± 3.1 million | 15.1 ± 2.8 million |

### Supplemental figures

**Fig S1.**
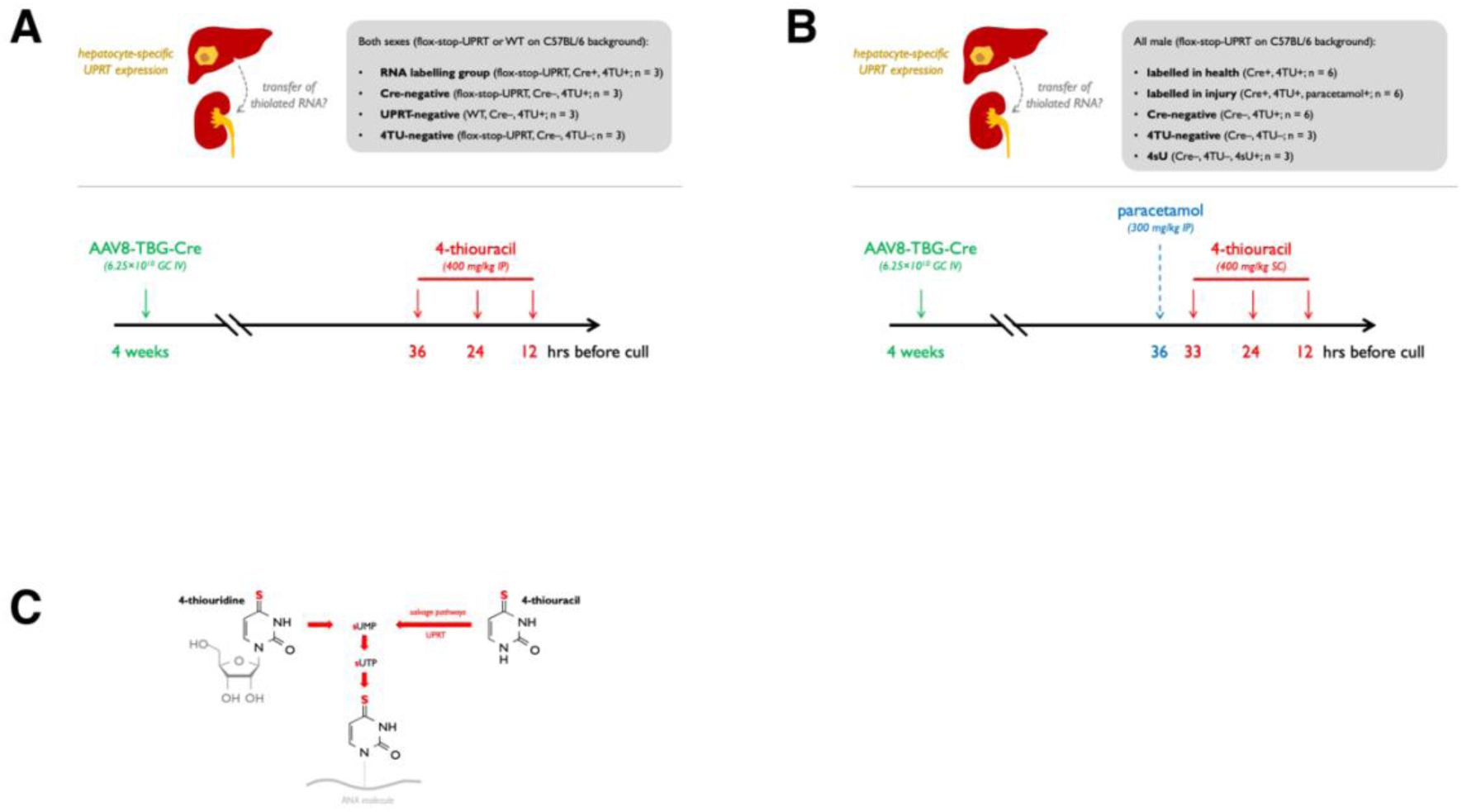
Experimental design. **A)** *Protocol for small initial experiment.* An AAV8 vector was used to deliver Cre recombinase specifically to hepatocytes in floxed-stop-UPRT mice approximately 4 weeks prior to RNA labelling. Mice were dosed with 4-thiouracil (400 mg per kg) at 36, 24 and 12 hrs prior to cull. **B)** *Liver injury protocol.* Mice in the injury group were dosed with paracetamol (300 mg per kg) at 36 hrs prior to cull; the first dose of 4-thiouracil was then given 3 hrs after paracetamol (i.e. 33 hrs prior to cull) and then at 24 and 12 hrs prior to cull. **C)** *Metabolism of 4TU and 4sU.* The ribonucleoside 4sU, 4-thiouridine, is imported into all metazoan cells by equilibrative nucleoside transporters and incorporated into nascent RNA. 4-thiouracil, 4TU, must first be converted into thiolated uridine monophosphate (sUMP) before it can be incorporated into RNA. In higher eukaryotes, salvage pathways converting 4TU to sUMP operate with very low efficiency. The expression of recombinant protozoan UPRT permits the efficient conversion of 4TU to sUMP and thence incorporation into RNA molecules.

**Fig S2.**
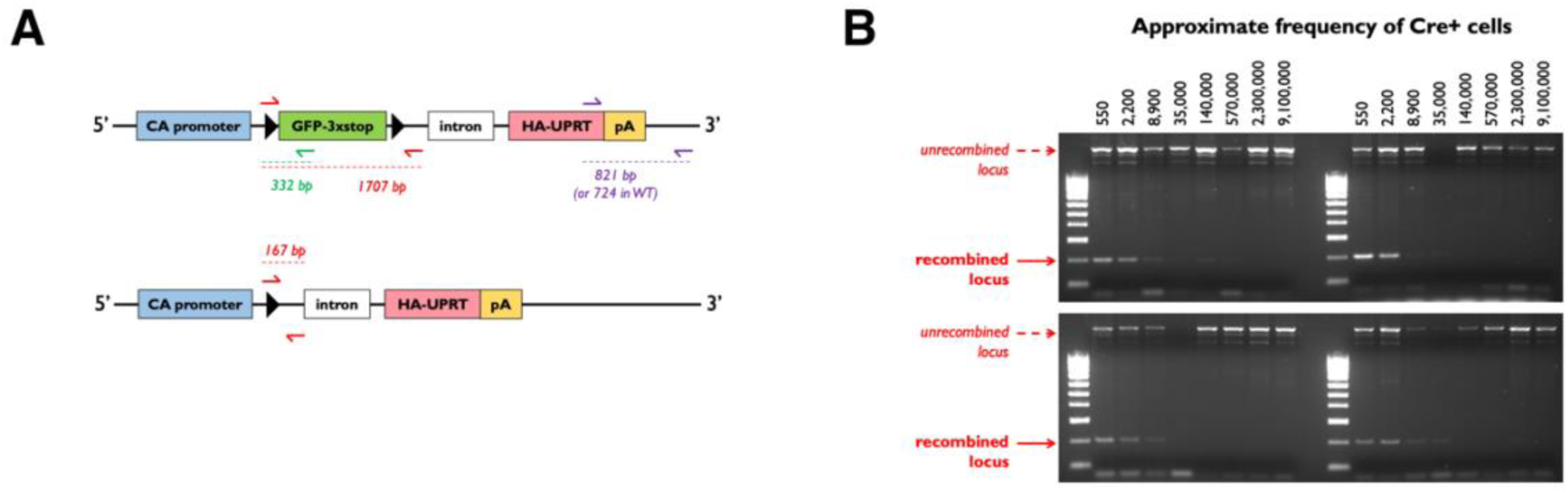
Genomic DNA (gDNA) recombination assay. **A)** *Design of recombination assay.* Primers (red) were designed to amplify a 1707 bp region of the unrecombined flox-stop-UPRT locus or a 167 bp region of the recombined locus. The green reverse primer was used to amplify a 332 bp region only from the unrecombined locus and purple primer set for genotyping. **B)** *Recombination dilution series in podo-UPRT.* gDNA was prepared from kidneys in podocin-UPRT mice and then used in a dilution series to estimate the sensitivity of the recombination assay. gDNA was diluted serially into gDNA from wild-type kidney and then used as a template in the PCR recombination assay reaction (n = 4). Assuming that podocytes constitute ∼3% of all kidney cells and that Cre recombination in podocytes is 100% efficient, the approximate frequency of recombined cells is shown at the top of the figure. The assay was reliably able to detect Cre recombination, even when the recombined cells were estimated to constitute ∼1 in 35,000 kidney cells and could detect recombination in some replicates down to an estimated frequency of 1 in 570,000 cells.

**Fig S3.**
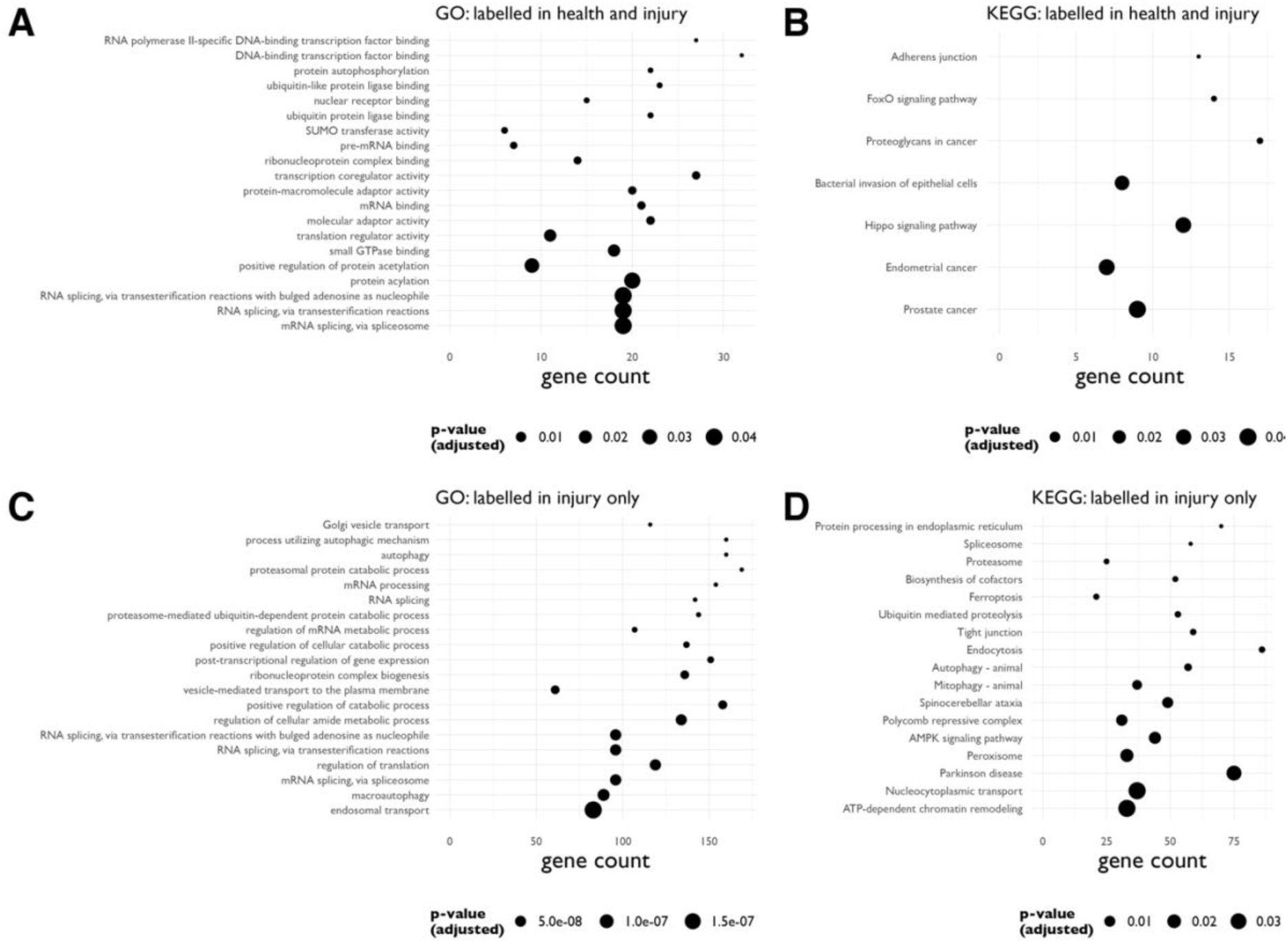
Pathway analyses of mRNAs likely to be transferred from liver to kidney. **A&B)** Gene ontology (**A**) and KEGG pathways (**B**) analyses of those mRNAs that were labelled in the kidney in both health and liver injury in 4TU-treated mice, but not in 4sU-treated mice (i.e. the set containing 478 mRNAs in **Error! Reference source not found.**E). **C&D**) GO (**C**) and KEGG (**D**) analyses of those mRNAs that were labelled in the kidney exclusively in liver injury (i.e. the set containing 3690 mRNAs in **Error! Reference source not found.**E). In these analyses, the set of labelled mRNAs was compared to a “background” set comprising all of the mRNAs present in the corresponding kidney RNAseq dataset.

